# Evidence for associations between Rey-Osterrieth Complex Figure test and motor skill learning in older adults

**DOI:** 10.1101/2020.09.27.315168

**Authors:** Jennapher Lingo VanGilder, Keith R. Lohse, Kevin Duff, Peiyuan Wang, Sydney Y. Schaefer

## Abstract

Age-related declines in motor learning may be related to poor visuospatial function. Thus, visuospatial testing could evaluate older adults’ potential for motor learning, which has implications for geriatric motor rehabilitation. To this end, the purpose of this study was to identify which visuospatial test is most predictive of motor learning within older adults. Forty-five nondemented older adults completed six standardized visuospatial tests, followed by three weekly practice sessions on a functional upper-extremity motor task. Participants were re-tested one month later on the trained task and another untrained upper-extremity motor task to evaluate the durability and generalizability of motor learning, respectively. Principal component analysis first reduced the dimensions of the visuospatial battery to two principal components for inclusion in a mixed-effects model that assessed one-month follow-up performance as a function of baseline performance and the principal components. Of the two components, only one was related to one-month follow-up. Factor loadings and post hoc analyses suggested that of the six visuospatial tests, the Rey-Osterrieth test (visual construction and memory) was related to one-month follow-up of the trained and untrained tasks. Thus, it may be plausible that older adults’ long-term motor learning capacity could be evaluated using the Rey-Osterrieth test, which would be feasible to administer prior to motor rehabilitation to indicate risk of non-responsiveness to therapy.

## 1. Introduction

Motor rehabilitation is important for recovering lost motor function and/or training compensatory movement patterns central for activities of daily life in older adults. While the capacity to learn and generalize motor skills is fundamental to the rehabilitative process, numerous studies indicate that aging negatively impacts motor learning (i.e., the extent to which one can achieve relatively-permanent changes in motor performance due to repeated practice (Schmidt, Lee, Winstein, Wulf, & Zelaznik, 2018)), such that older adults tend to learn and adapt movements to a reduced degree compared to younger adults (Raz, Williamson, Gunning-Dixon, Head, & Acker, 2000; Swinnen, 1998; Verwey, 2010; Wang, Williams, & Wilmut, 2020). As such, age-related motor learning impairments may in part explain why older adults tend to be less responsive to motor therapy than younger adults (e.g., Dobkin et al., 2014; Schaefer et al., 2019)).

A recent series of experimental studies has suggested that age-related declines in motor learning may be specifically associated with reduced visuospatial function (Bo, Borza, & Seidler, 2009; Langan & Seidler, 2011; Lingo VanGilder, Hengge, Duff, & Schaefer, 2018; Lingo VanGilder, Walter, Hengge, & Schaefer, 2019); *visuospatial function* refers to the broad spatial processes related to high-level vision and visuomotor integration (Jagaroo, 2009) and tends to decline across the adult lifespan sooner and at a faster rate than other cognitive functions (Murre, Janssen, Rouw, & Meeter, 2013). Indeed, visuospatial cognition may uniquely predict motor learning capacity in older adults, while the function of other cognitive domains (such as attention, language, delayed memory, etc.) may not (Lingo VanGilder et al., 2018; Toglia, Fitzgerald, O’Dell, Mastrogiovanni, & Lin, 2011). These findings build on the seminal work of Fleishman and Rich (1963), who observed that early improvements in visuomotor task performance were dependent upon visuospatial perception in a group of healthy young adult men. More recent work highlights the role of visuospatial working memory in other forms of motor learning, namely sensorimotor adaptation (Langan & Seidler, 2011; Schaffert, Lee, Neill, & Bo, 2017) and sequence learning (Bo et al., 2009; Bo & Seidler, 2009; Chan, Wu, Liang, & Yan, 2015) in both young and older adults. Similarly, an analogous effect was reported from experimental paradigms that trained participants on select activities of daily living, whereby the extent of improvement was related to visuospatial/executive function in stroke survivors (Toglia et al., 2011) and nondemented older adults (Lingo VanGilder et al., 2019).

It is therefore plausible that visuospatial assessment may have clinical utility in predicting responsiveness to motor therapy as a means to identify individuals who may need more intensive or targeted training. However, despite the myriad of studies supporting the functional relationship between visuospatial and motor cognition in nondemented older adults (e.g., Emerson et al., 2012), there are critical knowledge gaps that preclude translation of these findings into the clinic. First, as the visuospatial domain broadly encompasses cognitive constructs of perception, memory, construction, among others (described in further detail below), and previous work evaluated disparate aspects of visuospatial cognition, it remains unclear which visuospatial construct(s) is most predictive of motor learning ability in older adults. Second, previous studies relating visuospatial function and motor learning have not consistently used any given visuospatial test, and some have even used unvalidated, experimenter-derived assessments to attempt to quantify visuospatial function. No studies to date have directly compared how well one visuospatial assessment predicts motor learning relative to another. Thus, there is currently no clear candidate for which visuospatial assessment is best, despite there being a number of standardized visuospatial tests that may have merit (see Methods). Third, these previous studies measured motor learning as the amount of motor skill learned within one session or retained over consolidation periods of one day to one week, yet longer retention periods are arguably required based on the widely-accepted definition of ‘motor learning’ involving relatively permanent changes in performance due to experience (Schmidt et al., 2018).

To address these knowledge gaps, the present study was designed to directly compare multiple standardized visuospatial tests to identify which was most predictive of long-term motor learning outcomes (i.e., skill retention and transfer) in older adults. Based on previous work from our lab and others’, we hypothesized that tests of visuospatial perception and memory would positively correlate with the degree of motor skill retained and transferred one month after extensive training. Results of this study will advance our understanding of human visuospatial and motor cognition, and serve as a critical step towards implementing prognostic testing within motor therapy treatment plans.

## 2. Methods

All experimental procedures were approved by Arizona State University’s Institutional Review Board prior to participant recruitment. Forty-five right-handed participants (28 female) with a mean ± SD age of 70.38 ± 6.77 years with no self-reported history of a previous neurological or psychiatric condition (e.g., stroke, schizophrenia, etc.) participated in this study. Participants completed a series of examinations to characterize sensory, motor, daily functioning, and mood. Bilateral index finger sensation (Semmes-Weinstein monofilaments) and grip strength (hand dynamometer (Jamar Technologies)) were collected. Participant handedness was determined using the Edinburgh Questionnaire (Oldfield, 1971) and dexterity of the nondominant hand was evaluated using the Grooved Pegboard test. Participants also completed the Activities of Daily Living Questionnaire (Katz, Downs, Cash, & Grotz, 1970) to determine the extent they independently perform six common physical activities, where the best possible score of 6 indicates full independence. The Geriatric Depression Scale was completed at the beginning of each visit. Participant data for all measures are provided in Table 1.

**Table 1.**
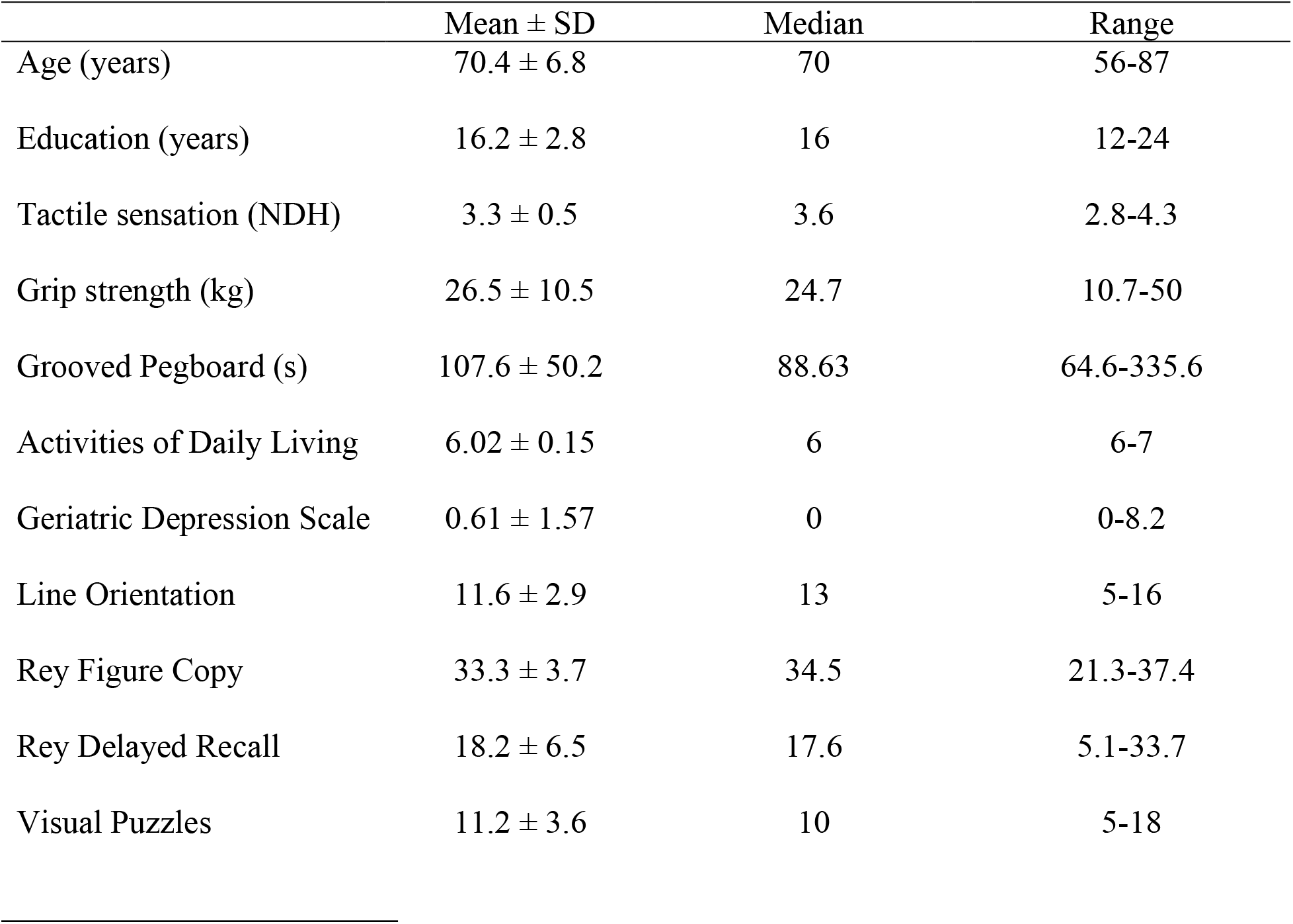
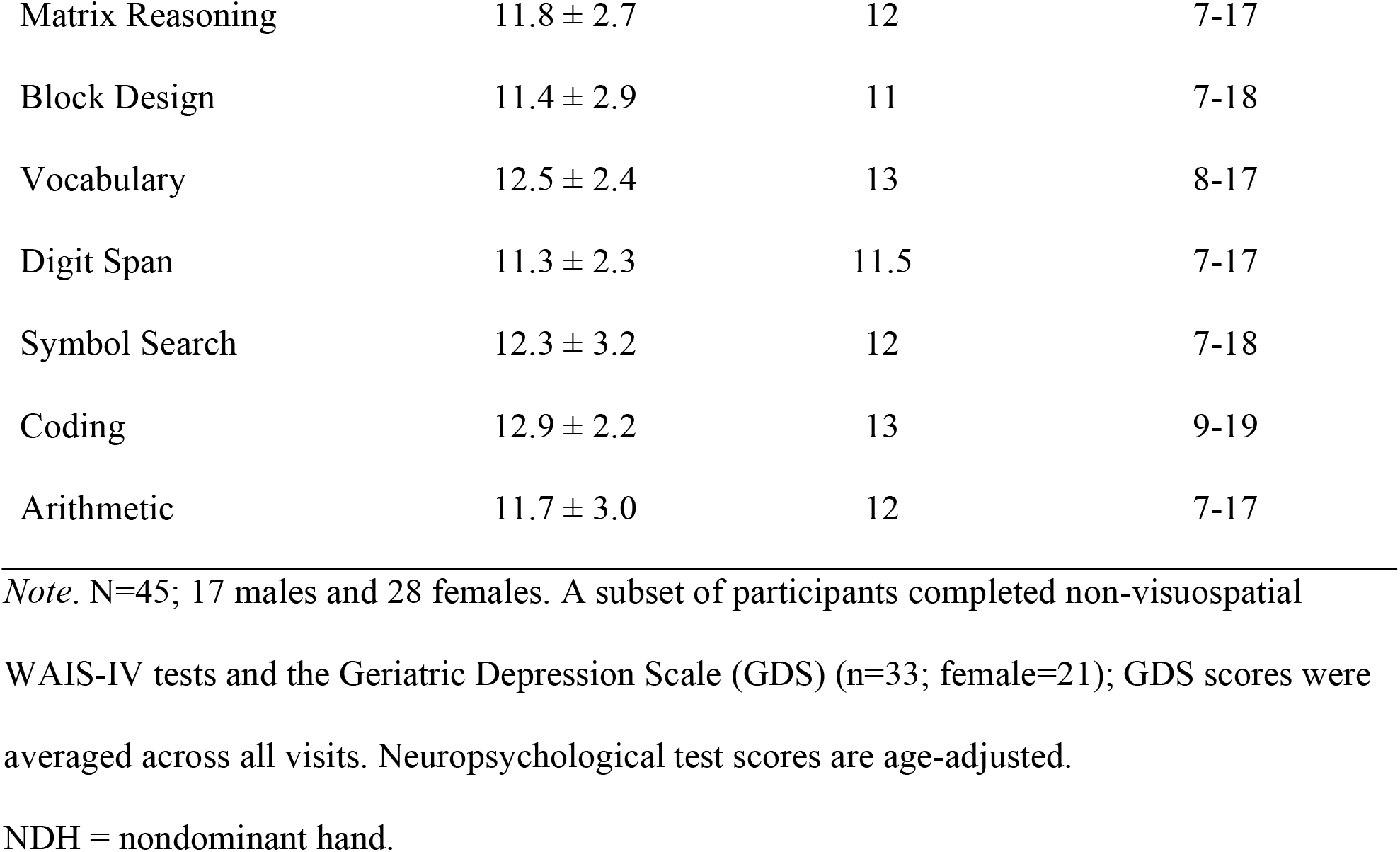
Participant characteristics and neuropsychological data.

| | Mean ± SD | Median | Range |
| --- | --- | --- | --- |
| Age (years) | 70.4 ± 6.8 | 70 | 56-87 |
| Education (years) | 16.2 ± 2.8 | 16 | 12-24 |
| Tactile sensation (NDH) | 3.3 ± 0.5 | 3.6 | 2.8-4.3 |
| Grip strength (kg) | 26.5 ± 10.5 | 24.7 | 10.7-50 |
| Grooved Pegboard (s) | 107.6 ± 50.2 | 88.63 | 64.6-335.6 |
| Activities of Daily Living | 6.02 ± 0.15 | 6 | 6-7 |
| Geriatric Depression Scale | 0.61 ± 1.57 | 0 | 0-8.2 |
| Line Orientation | 11.6 ± 2.9 | 13 | 5-16 |
| Rey Figure Copy | 33.3 ± 3.7 | 34.5 | 21.3-37.4 |
| Rey Delayed Recall | 18.2 ± 6.5 | 17.6 | 5.1-33.7 |
| Visual Puzzles | 11.2 ± 3.6 | 10 | 5-18 |
| Matrix Reasoning | 11.8 ± 2.7 | 12 | 7-17 |
| Block Design | 11.4 ± 2.9 | 11 | 7-18 |
| Vocabulary | 12.5 ± 2.4 | 13 | 8-17 |
| Digit Span | 11.3 ± 2.3 | 11.5 | 7-17 |
| Symbol Search | 12.3 ± 3.2 | 12 | 7-18 |
| Coding | 12.9 ± 2.2 | 13 | 9-19 |
| Arithmetic | 11.7 ± 3.0 | 12 | 7-17 |

*Note.* N=45; 17 males and 28 females. A subset of participants completed non-visuospatial
WAIS-IV tests and the Geriatric Depression Scale (GDS) (n=33; female=21); GDS scores were averaged across all visits. Neuropsychological test scores are age-adjusted.
NDH = nondominant hand.

### 2.1. Visuospatial and other cognitive tests

To address the purpose of this study, participants completed a battery of standardized cognitive tests that were selected in consultation with a licensed clinical neuropsychologist (KD) prior to the start of the study to comprehensively evaluate the visuospatial domain:

- Benton Judgment of Line Orientation (Benton et al., 1994): This 30-item test is a validated measure of visual perception in which the participant selects a line from an array that is in the same direction and orientation as a test line shown above the array. Correct answers receive points, with a maximum score of 30 points.
- Rey-Osterrieth Complex Figure Test (Osterrieth, 1944): The Copy trial is a validated measure of visual construction in which the participant copies a complex image on a separate sheet of paper as accurately as possible. Correctly copied elements receive points, with a maximum score of 36 points. The Delayed Recall trial is a validated measure of visual memory in which the participant is asked to redraw the complex image from memory after a 30-minute period. Scores are identical to the Copy trial.
- Visual Puzzles: This 26-item validated measure of visual reasoning within the Wechsler Adult Intelligence Scale, 4^th^ edition (Wechsler, 1955) (WAIS-IV). The participant is presented with a visual design and asked to select three constituent images that, when combined, reconstruct the completed puzzle. Correct answers receive points, with a maximum score of 26 points.
- Matrix Reasoning: This 26-item validated measure of visual abstract-problem solving, also within the WAIS-IV. The participant is presented with an incomplete visual matrix or series of images and is asked to select an option that completes the matrix or series. Correct answers receive points, with a maximum score of 26 points.
- Block Design: This 14-item validated measure of visual (object) construction, also within the WAIS-IV. The participant is presented with a 2- or 3-dimensional model and is asked to use red-and-white blocks to recreate the model. Correct recreations receive points, with bonus points for quicker responses, with a maximum score of 48 points.

It is noted that these tests may also require executive function or multiple aspects of visuospatial cognition (as is the case for Rey Complex Figure Copy, in which performance relies on planning and organization (executive function) as well as visuospatial construction and perception (Fillit, Rockwood, & Young, 2016). Again, the purpose of this study is to identify the clinical visuospatial test most predictive of one-month motor learning outcomes in older adults, and *not* to isolate the cognitive mechanism (i.e., which constructs) underlying motor learning. Additional WAIS-IV tests were used to exclude and characterize language development (Vocabulary), processing speed (Coding and Symbol Search), working memory (Arithmetic), and auditory attention (Digit Span); any score ≤ 5^th^ percentile was considered a cutoff for study exclusion to minimize the likelihood of enrolling participants with dementia. These WAIS-IV test scores were not evaluated as predictors of motor learning, however, based on their lack of association in previous studies. All raw neuropsychological test scores were age-adjusted according to the test instructions (e.g., WAIS-IV) or published methods (Caffarra, Vezzadini, Dieci, Zonato, & Venneri, 2002; Ivnik, Malec, Smith, Tangalos, & Petersen, 1996). In all cases, higher scores indicated better performance on these cognitive tests.

### 2.2. Motor tasks

After completing all sensorimotor and cognitive assessments, baseline performance was collected for participants on two motor tasks designed to mimic important activities of daily living: functional reaching and functional dexterity. (Justification of the selection of these tasks are provided below in section 2.3.) The functional reaching task involved upper extremity movements similar to self-feeding (i.e., using a spoon to acquire and transport objects from one location to another), whereas the functional dexterity task involved upper extremity movements necessary for self-dressing (i.e., fastening buttons). For the functional reaching task, participants used a conventional plastic spoon with their nondominant hand to acquire and transport two raw pinto beans at a time from a central cup (centered at their midline) to one of three target cups located radially about the center cup at a distance of 16 cm. All cups had a 9.5 cm diameter and were 5.8 cm tall. Participants first reached towards the ipsilateral cup, then the middle cup, and then the contralateral cup; this sequence was repeated until the last two beans from the center cup were deposited into the last target cup. Trial time began when the participant picked up the spoon, which was located next to the cup. The measure of performance on this task was trial time (i.e., the time it took to complete 15 reaches), with lower values indicating better performance. The functional dexterity task involved a wooden board (61 cm x 34 cm) with a piece of heavyweight linen fabric adhered to the back; the fabric folded around the front of the board to form a placket down its center. One side of the fabric contained 10 one-inch buttons sewn along its edge, 5.3 cm apart, while the opposing piece contained 10 complementary buttonholes. The board was placed at the participant’s midline at the beginning of each trial. Participants were instructed to use their nondominant hand to fasten the buttons sequentially as fast as possible. Trial time began when the participant flipped over the button-side piece of fabric and ended when the last button was fastened. Again, lower trial times indicate better performance. Schematics for each task are provided in Figure 1, while additional details regarding experimental apparatus and administration (for both motor tasks) have been previously published (Lingo VanGilder et al., 2019; Schaefer & Hengge, 2016) and are publicly available on Open Science Framework (https://osf.io/phs57/wiki/Functional_reaching_task/).

**Figure 1.**
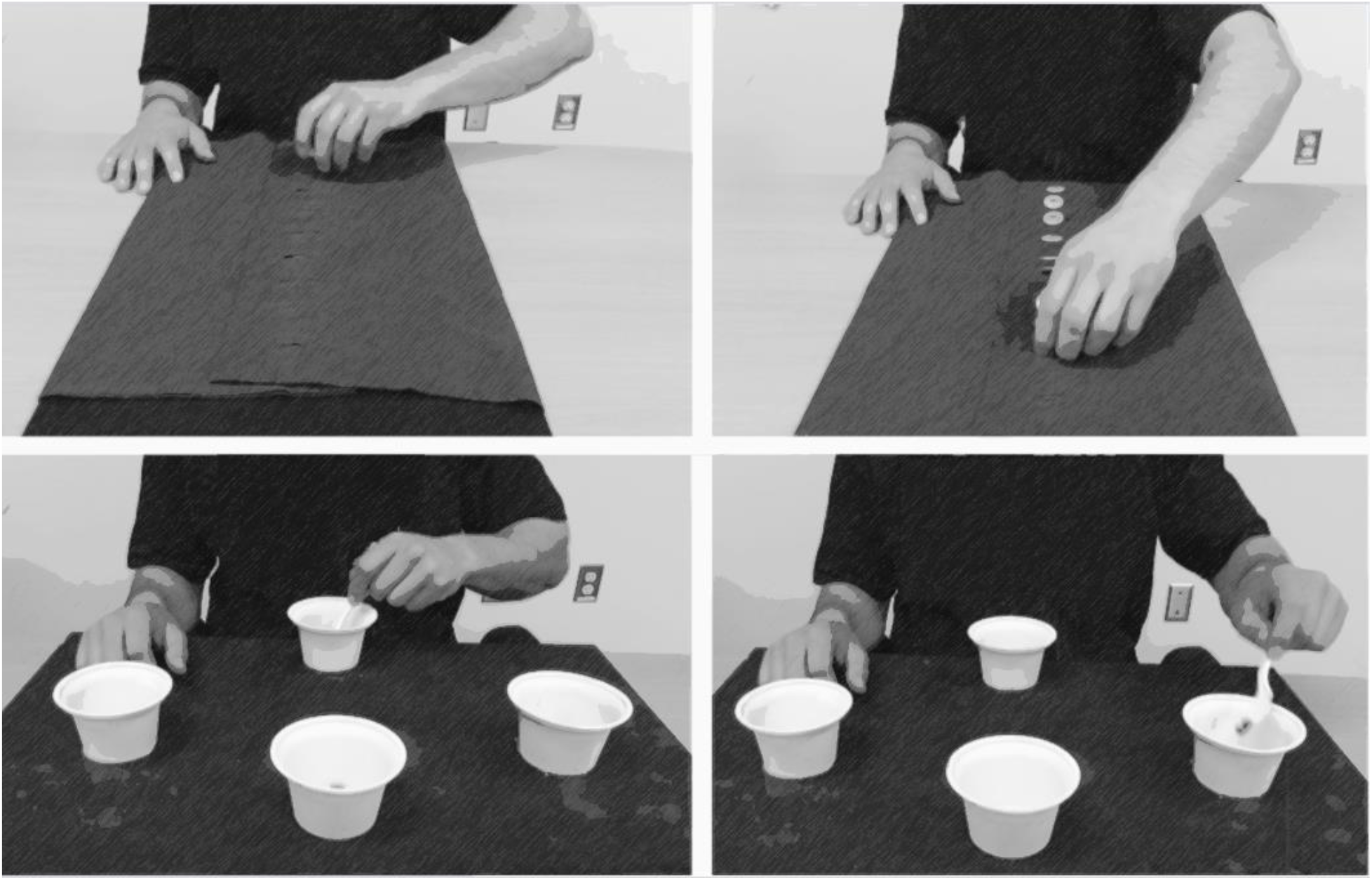
Participants used their nondominant hand to complete two functional motor tasks: A functional dexterity task (top panel) and a functional reaching task (bottom panel). “Dexterity and Reaching Motor Tasks” by MRL Laboratory is licensed under CC BY 2.0.

### 2.3 Motor training protocol

On Day 1, participants were evaluated on all sensory, motor, and cognitive (including all visuospatial) assessments, and baseline performance of the functional upper extremity motor tasks. On Days 8, 15, and 22 (i.e., one, two, and three weeks after baseline), participants completed 50 training trials of just the functional reaching task, thereby simulating task-specific training. Participants received no training on the functional dexterity task, which was used as the transfer task in this study. Thus, participants underwent three 50-trial functional reaching training sessions, one week apart. One month after the last training session, participants were re-tested on both the functional reaching and functional dexterity tasks to evaluate long-term skill retention and transfer, respectively. These particular motor tasks were selected for this study because this training paradigm has previously shown transfer of learning in both stroke (Schaefer, Patterson, & Lang, 2013) and cognitively-intact adult (Lingo VanGilder et al., 2019; Schaefer & Lang, 2012) populations. In other words, there is previous evidence that transfer occurs from the functional reaching task to the functional dexterity task. Furthermore, the dose and timing of motor training was based on previous work demonstrating their efficacy in promoting lasting training effects (retention and transfer) (e.g., Schaefer et al., 2015; Walter et al., 2019). The training in this study was *not* intended to mimic the delivery of motor therapy in standard care (as the dose of training here exceeds that of clinical motor therapy) (Kimberley, Samargia, Moore, Shakya, & Lang, 2010; Lang, MacDonald, & Gnip, 2007; Lang et al., 2009).

### 2.4 Statistical analysis

Statistical analyses were performed using JMP Pro 13.1 software (SAS Institute Inc., Cary, NC) and R 3.6.1 (R Core Team). Wilcoxon signed-ranked tests were used to first verify that there was a significant amount of improvement in the trained functional reaching (i.e., retention) and untrained dexterity (i.e., transfer) tasks from baseline to one-month follow-up. Because visuospatial tests do not evaluate a unitary visual construct (e.g., a test may be a validated measure of visual construction, yet performance on that test may also engage other aspects of visuospatial cognition, such as perception, reasoning, etc.), it is likely that the visuospatial tests used in this study may overlap in the visuospatial constructs they test (i.e., they are not independent of each other). To account for collinearity among these visuospatial tests, all age-adjusted visuospatial scores (Benton Judgment of Line Orientation, Rey Complex Figure Copy and Delayed Recall, Visual Puzzles, Matrix Reasoning, and Block Design) were subjected to an *a priori* principal component analysis to reduce dimensionality for inclusion in regression analysis. This step serves two important purposes: (1) it isolates the shared variance between tests that would be excluded if all of the visuospatial scores were put into the same multivariable regression in parallel, and (2) it reduces the number of statistical tests that would need to be run if each visuospatial score was tested in serial (i.e., six different statistical tests, one for each visuospatial measure), thereby controlling for Type-I error. Loading matrices were used to identify which visuospatial tests loaded on to which principal component (PC). The Kaiser-Meyer-Olkin measure of sampling adequacy was calculated to confirm our sample size provided appropriate stability for principal component analysis (Kaiser, 1974).

All PCs with Eigenvalues > 1.00 were then carried forward to a mixed-effects regression model that also used baseline performance and task (reaching vs. dexterity) as predictor variables of one-month follow-up performance. As we had six candidate visuospatial tests to consider rather than a single ‘gold standard’ visuospatial test, a family-wise error correction could have been imposed, but being cognizant of the cost of Type-II errors in this exploratory study was also important; we reasoned Type-II errors to be more costly. ^1^

Mixed-effects regression models included factors of task, baseline performance, and all PCs. To follow-up statistically interesting effects of PCs (or Task x PC interactions), the age-adjusted scores from the tests that loaded on that PC were then entered back into the mixed-effects model in place of the PC, one test at a time. This post hoc analysis determined if an individual test (i.e., Line Orientation, Figure Copy, etc.) could in fact be used to predict skill retention or transfer.

## 3. Results

Participant characteristics are provided in Table 1.

Results from the Wilcoxon signed-rank tests indicated that participants significantly improved their performance from baseline to one-month follow-up on both motor tasks. For the functional reaching task, participants improved by a mean of 14.7 seconds from baseline to follow-up (Z = - 261, p < 0.0001), and for the functional dexterity task, participants improved by 6.6 seconds despite no training (Z = −110, p < 0.018). These values are reported simply to confirm that both tasks showed significant improvement; given that the tasks themselves are functionally quite different (Schaefer et al., 2013), the magnitude of improvement for one task is not meant to be compared to that for the other task. Mean values for baseline and one-month follow-up across participants are shown in Figure 2 for both tasks, as well as the overall training data (Fig. 2). This demonstrates the feasibility and efficacy of the motor training paradigm in this sample.

**Figure 2.**
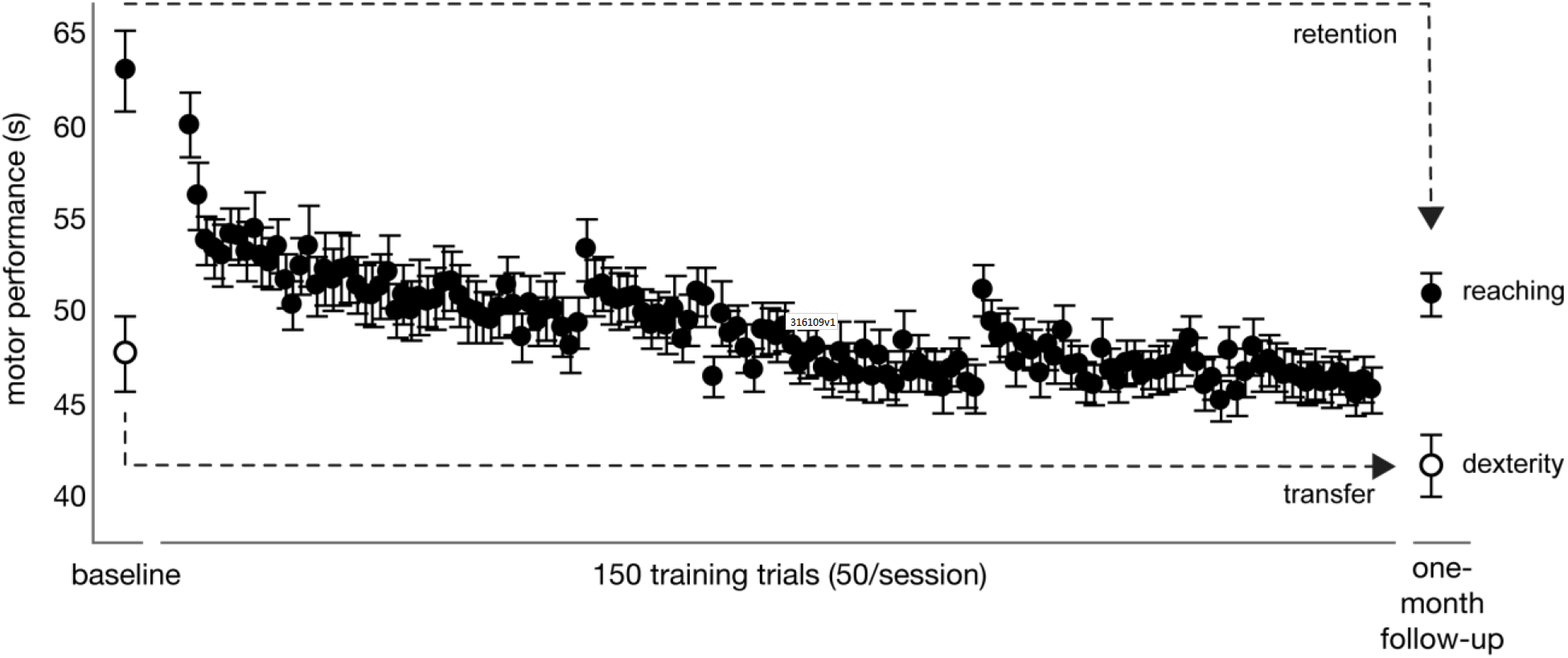
Participants completed a baseline trial of the reaching and dexterity motor tasks, then completed 50 training trials on the reaching task during three weekly sessions (totaling 150 trials). Participants were retested on the trained (reaching) and untrained (dexterity) task one month later to determine skill retention and transfer, respectively. Mean motor performance (trial time in seconds) is plotted on the y-axis, where lower values indicate better performance. • = trained reaching task; ∘ = untrained dexterity task. Error bars indicate standard error.

Principal component analysis was used to reduce the dimensionality across the age-adjusted scores from the six visuospatial tests and account for shared variance among them. To confirm our sample size provided appropriate stability for principal component analysis, we computed the measure of sampling adequacy (MSA = 0.77), which indicates the sample size was reasonable and our component loadings would be moderately stable. Two PCs emerged with Eigenvalues > 1.00, and when combined accounted for 69.7% of the variance. The visuospatial tests and corresponding factor loadings for each PC are provided in Table 2. Results indicated that Block Design and Matrix Reasoning primarily loaded on PC1, whereas Rey-Osterrieth Complex Figure Copy and Delayed Recall primarily loaded on PC2.

**Table 2.**
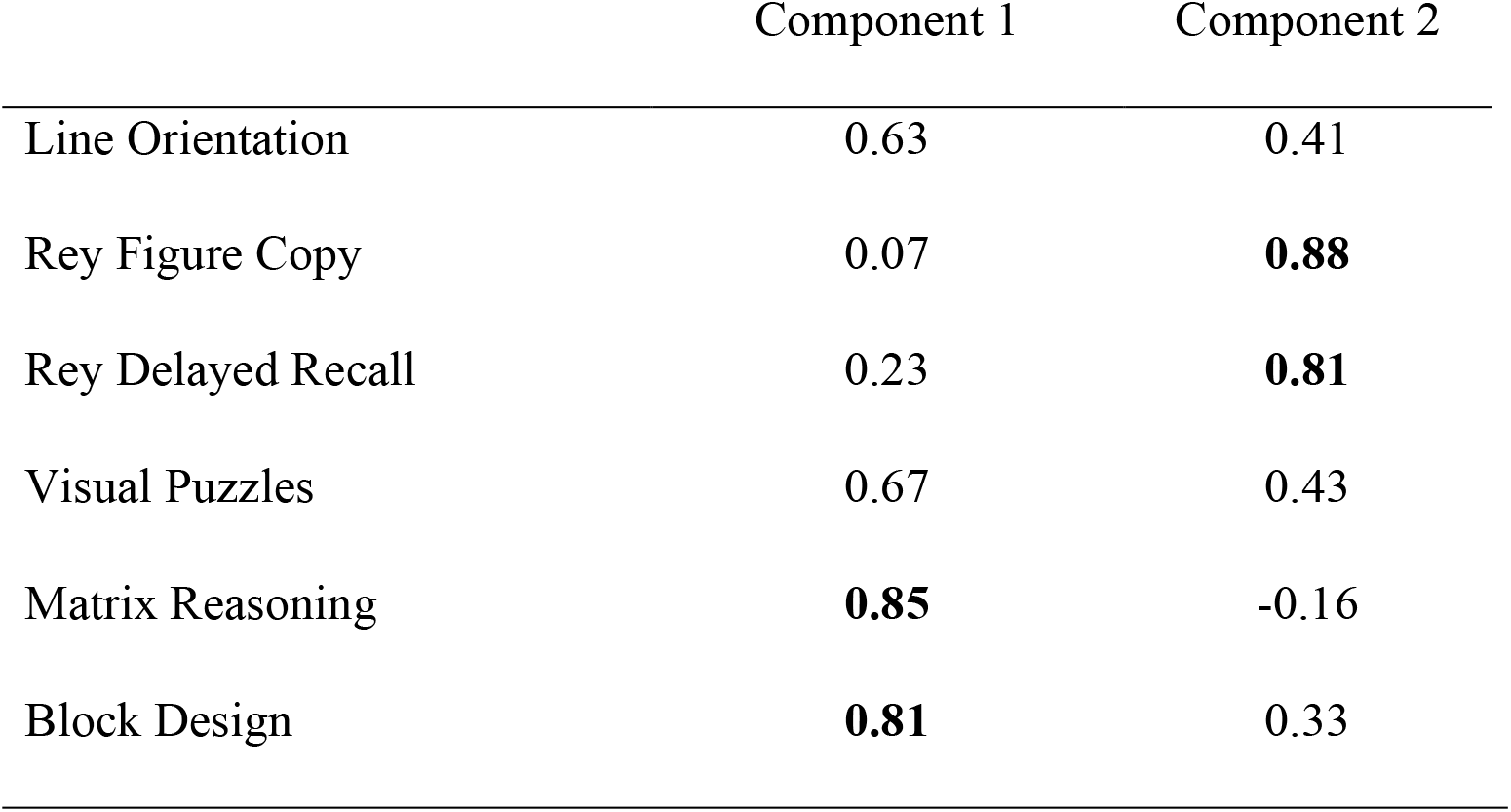
Factor loadings for principal components.

Additional bivariate scatterplots between participant performance on each visuospatial test are provided in the Supplementary Material (eTable 1) and support the above PCA findings.

To determine which PC(s) predicted motor skill retention and transfer, PC1 and PC2 were entered in a mixed-effects model as predictors of one-month follow-up motor performance, along with baseline motor performance, a ‘task’ factor (functional reaching vs. functional dexterity), and task interactions (see Table 3).

**Table 3.** Parameters from the mixed-effect regression model explaining one-month follow-up performance.

| Random-Effects |  |  |
| --- | --- | --- |
| *Name* | *Variance* | *SD* |
| Participant | 1.70 | 1.31 |

Residual 38.76 6.23 Based on 90 observations from 45 participants.
| Fixed-Effects |  |  |  |  |  |
| --- | --- | --- | --- | --- | --- |
| *Name* | *Estimate ($\beta$)* | *SE* | *df* | *t-value* | *p-value* |
| Intercept | 46.96 | 0.78 | 60.02 | 59.85 | <0.001* |
| Baseline | 0.49 | 0.05 | 78.14 | 9.62 | <0.001* |
| Task | -0.88 | 0.76 | 60.82 | -1.16 | 0.251 |
| PC1 | -0.09 | 0.40 | 43.45 | -0.22 | 0.828 |
| PC2 | -1.08 | 0.66 | 44.17 | -1.63 | 0.110 |
| Baseline x Task | 0.09 | 0.05 | 73.97 | 1.76 | 0.083 |
| PC1 x Task | -0.07 | 0.39 | 43.45 | -0.19 | 0.850 |
| PC2 x Task | -1.27 | 0.64 | 44.13 | -1.99 | 0.053 |

*Note.* Baseline was mean-centered and Task was contrast coded with Untrained = +1 and Trained = -1. Degrees of freedom were estimated based on the Welch-Satterthwaite approximation (Satterthwaite, 1946; Welch, 1947). Further, a covariate interaction was controlled for to get the most unbalanced estimate of the interaction of interest (Yzerbyt, Muller, & Judd, 2004). PC = principal component.

As expected, regression analysis revealed a statistically significant relationship between baseline and one-month follow-up performances for both motor tasks (p < 0.001). This relationship was slightly stronger for the untrained dexterity task than the trained reaching task, but the Baseline × Task interaction was not statistically significant (p = 0.083). There was not a significant main effect of PC1 (p = 0.83), nor was there a significant PC1 × Task interaction (p = 0.85), suggesting that PC1 (i.e., the shared variance between tests that loaded on this PC) has a minimal relationship with one-month follow-up performance and that this relationship does not change as a function of task. While there was not a significant main effect of PC2 (p = 0.11), there was evidence for a PC2 × Task interaction (p = 0.053), suggesting PC2 (namely, the Rey-Osterrieth test) may be related to follow-up performance, particularly for the untrained task (i.e., skill transfer).

When considering 95% confidence intervals for each effect rather than their statistical significance in isolation (Wasserstein, Schirm, & Lazar, 2019), there was again little evidence to suggest that PC1 was related to one-month follow-up performance (main-effect β = −0.09, CI = [−0.90, 0.73]), or changed as a function of task (interaction β = −0.07, CI = [−0.85, 0.71]). In contrast, PC2 appeared to have a stronger relationship with one-month follow-up performance overall (main-effect β = −1.08, CI = [−2.41, 0.25]), which was potentially stronger for the untrained task (i.e., skill transfer) compared to the trained task (i.e., skill retention) (interaction β= −1.27, CI = [−2.55, 0.01]). Given the history of past work showing that (various) visuospatial tests correlate with individual differences in motor learning outcomes, we decided to investigate this interaction further.

Because the Rey-Osterrieth Complex Figure test primarily loaded on to PC2, and there was evidence that PC2 was related to motor skill transfer, additional analyses evaluated whether this relationship was predicated upon the performance of the Figure Copy or Delayed Recall trial. Individual mixed-effects models analogous to those shown in Table 3 were independently conducted for the Figure Copy and Delayed Recall trials of the Rey-Osterrieth Test, in place of PC2 (all other variables remained in the model) (Tables 4 and 5, respectively).

**Table 4.** Parameters from the mixed-effect regression model using Figure Copy to explain one-month follow-up performance.

| Random-Effects |  |  |  |  |  |
| --- | --- | --- | --- | --- | --- |
| *Name* | *Variance* | *SD* |  |  |  |
| Participant | 3.07 | 1.75 |  |  |  |
| Residual | 37.67 | 6.14 |  |  |  |
| Based on 90 observations from 45 subjects. |  |  |  |  |  |
| Fixed-Effects |  |  |  |  |  |
| *Name* | *Estimate ($\beta$)* | *SE* | *df* | *t-value* | *p-value* |
| Intercept | 27.34 | 9.35 | 51.56 | 2.92 | 0.005* |
| Baseline | 0.49 | 0.05 | 80.45 | 9.55 | <0.001* |
| Task | -5.11 | 2.88 | 72.25 | -1.77 | 0.081 |
| PC1 | 0.23 | 0.51 | 41.38 | 0.45 | 0.657 |
| FC | -0.23 | 0.25 | 43.72 | -0.94 | 0.352 |
| Baseline x Task | 0.08 | 0.05 | 72.90 | 1.53 | 0.130 |
| PC1 x Task | 0.62 | 0.47 | 41.39 | 1.31 | 0.199 |
| FC x Task | -0.54 | 0.23 | 43.58 | -2.35 | 0.023* |

*Note.* Baseline was mean-centered and Task was contrast coded with Untrained = +1 and Trained = -1. Degrees of freedom were estimated based on the Welch-Satterthwaite approximation.
PC1 = Principal component 1; FC = Figure Copy.

**Table 5.** Parameters from the mixed-effect regression model using Delayed Recall to explain one-month follow-up performance.

| Random-Effects |  |  |  |  |  |
| --- | --- | --- | --- | --- | --- |
| *Name* | *Variance* | *SD* |  |  |  |
| Participant | 0.00 | 0.00 |  |  |  |
| Residual | 40.97 | 6.40 |  |  |  |
| Based on 90 observations from 45 subjects. |  |  |  |  |  |
| Fixed-Effects |  |  |  |  |  |
| *Name* | *Estimate ($\beta$)* | *SE* | *df* | *t-value* | *p-value* |
| Intercept | 26.89 | 4.33 | 90.00 | 6.21 | <0.001* |
| Baseline | 0.48 | 0.05 | 90.00 | 9.55 | <0.001* |
| Task | -6.81 | 2.89 | 90.00 | -2.35 | 0.021* |
| PC1 | 0.79 | 0.54 | 90.00 | 1.47 | 0.146 |
| DR | -0.35 | 0.15 | 90.00 | -2.36 | 0.020* |
| Baseline x Task | 0.11 | 0.05 | 90.00 | 2.10 | 0.039* |
| PC1 x Task | 0.08 | 0.54 | 90.00 | 0.15 | 0.885 |
| DR x Task | -0.05 | 0.15 | 90.00 | -0.31 | 0.756 |

*Note.* Baseline was mean-centered and Task was contrast coded with Untrained = +1 and Trained = -1. Degrees of freedom were estimated based on the Welsh-Satterthwaite approximation.
PC1 = Principal component 1; DR = Delayed Recall.

Results indicated that Figure Copy scores were not reliably related to one-month follow-up performance overall (main-effect β = −0.23, CI = [−0.72, 0.26], p = 0.35), but showed a statistically more negative relationship to one-month follow-up performance on the untrained task (i.e., skill transfer) than that for the trained task (interaction β = −0.54, CI = [−0.07, 0.26], p = 0.02). Again, lower trial times correspond to better performance in our tasks; thus, results indicate that better Figure Copy scores predicted more skill transfer at one-month follow-up. Conversely, Delayed Recall scores were negatively related to performance overall (main-effect β = −0.35, CI = [−0.65, −0.05], p = 0.02) and this relationship was not statistically different between the trained and untrained tasks (interaction β = −0.05, CI = [−0.35, −0.25], p = 0.76), indicating higher Delayed Recall scores predicted better skill retention and transfer at one-month follow-up. Collectively, these findings suggest that of all the visuospatial tests administered in this study, the Rey-Osterrieth Complex Figure test is the most predictive of motor skill learning.

In the absence of an *a priori* power analysis, we used Monte Carlo simulation (Green & MacLeod, 2016) to test the sensitivity we achieved for the PC2 x Test interaction. These simulations assume that other aspects of the model are held constant (e.g., random-effects, variance explained by other parameters, etc.), but return the estimated power to detect a fixed effect at different magnitudes. At our sample size (N=45), we had only marginal power to actually detect what we observed (∼60%). We do have some confidence in this effect as it conceptually replicated past work, but future studies seeking to replicate this effect should use larger samples.

## 4. Discussion

The purpose of this study was to identify which visuospatial test(s) was most predictive of skill retention and transfer one-month after extensive motor training in nondemented older adults. Results indicated that older adults learned both functional upper extremity motor tasks (i.e., skill retention and transfer), and provided new evidence showing that among the visuospatial cognitive abilities evaluated, only construction and delayed visuospatial memory (i.e., Rey-Osterrieth Complex Figure test) predicted long-term learning. These findings are consistent with experimental studies that demonstrated figure drawing performance partially explained short-term skill transfer in normative aging (Lingo VanGilder et al., 2019) and individuals with stroke (Toglia et al., 2011). Overall, results support that visuospatial testing could be implemented in geriatric rehabilitation to estimate long-term motor learning potential.

### 4.1 Why might Rey-Osterrieth Complex Figure Tests be related to motor skill transfer?

A possible explanation for the observed relationship between transfer of motor learning and the Rey-Osterrieth test score is that the cognitive processes underlying these behaviors use a shared neural network. The strongest effect observed in our model comparison was Figure Copy performance (i.e., visuospatial construction), specifically. Visuospatial construction represents the ability to reconstruct a visual percept (e.g., an image or object) using its constituent parts (Mervis, Robinson, & Pani, 1999). Figure Copy performance is a function of graphomotor ability and may also rely on interplay between parallel cognitive networks underlying high-level motor control, visuomotor transformation, and multistep object use (Chen et al., 2016). Human clinical studies of dementia have shown that in addition to visuospatial construction, Figure Copy performance is also associated with multifaceted spatial constructs such as perception and working memory (Biesbroek et al., 2014; Freeman et al., 2000; Possin, Laluz, Alcantar, Miller, & Kramer, 2011). While the primary cognitive processes of skill transfer are less understood, it has been suggested that transfer of learning depends upon both short- and long-term memory. For instance, motor control theory suggests that transfer exploits abstract motor memories of the learned skill (Schmidt et al., 2018), and experimental studies indicate the degree of motor transfer in both young and healthy older adults is associated with spatial working memory (Langan & Seidler, 2011). It seems reasonable then, that Figure Copy performance may predict motor skill transfer as both behaviors rely on spatial working memory. It is noted, however, that although previous studies reported Figure Copy performance was related to working memory, Rey-Osterrieth Complex Figure Copy has not been validated for this purpose.

This theoretical framework is supported by neuroanatomical findings that indicate skill transfer and Rey-Osterrieth performance have similar structural and functional neural correlates as those with spatial memory. For example, functional and structural magnetic resonance imaging studies have implicated the role of the right inferior parietal lobule in the degree of skill transfer (Seidler, 2010), spatial working memory (Haley et al., 2008), and Figure Copy (Biesbroek et al., 2014). Moreover, cortical activations within the middle occipital gyri (Seidler, 2010) and dorsolateral prefrontal cortex (Kantak & Winstein, 2012) have been observed during both skill transfer and Figure Copy performance (Biesbroek et al., 2014; Possin et al., 2011). While collectively these findings suggest a common neural substrate may underlie skill transfer and visuospatial construction behavior, future work will examine the neuroanatomical correlates of this behavioral relationship within a homogeneous sample, and evaluate causal mechanisms (i.e., does the parietal, occipital, and/or frontal cortices mediate the relationship between transfer of motor learning and visuospatial construction ability?).

The Delayed Recall trial was also related to motor skill retention and transfer. Delayed Recall performance relies on the integration of perceptual-motor and planning-related executive abilities (González Viéitez, 2019), and has been used clinically to measure nonverbal episodic memory (Wong, Flanagan, Savage, Hodges, & Hornberger, 2014). Structural neuroimaging studies that evaluated episodic memory using Delayed Recall suggest that the mnemonic constructs required to reconstruct the figure primarily depend upon prefrontal networks (Wong et al., 2014), and that there may exist a left-hemispheric bias (Ostby, Tamnes, Fjell, & Walhovd, 2012). Indeed, a seminal neuroimaging study that evaluated visuospatial episodic memory recall in healthy adults observed increased cerebral blood flow to the left dorsolateral prefrontal cortex (among other regions) during task performance (Schacter et al., 1995). Consistent with these findings, experimental studies evaluating the structural neural correlates of long-term motor skill retention (e.g., a retention period > four weeks) reported increased cortical volume within primary motor and dorsoparietal (Sampaio-Baptista et al., 2014) and left prefrontal (Taubert et al., 2010) regions were positively correlated with greater amounts of skill retention. While collectively these studies may implicate left prefrontal networks in episodic and motor skill recall, the neural basis for the predictive relationship between Delayed Recall test scores and one-month skill retention remains unexplored.

### 4.2 Limitations and future directions

Although all training was completed with the nondominant (left) hand, the question emerges as to whether our findings are limited to right-hemispheric networks, or if they generalize to motor learning processes overall. While data from the current study do not directly resolve this issue, we hypothesize our results are generalizable based on previous work by Jeunet et al. (Jeunet, Jahanpour, & Lotte, 2016; Jeunet, Kaoua, Subramanian, Hachet, & Lotte, 2015). Visuospatial ability has been shown to predict learning of a motor-imagery task controlled by an electroencephalography-based brain-computer interface, a task that involves no effector (hand) at all. Further, visuospatial working memory has been associated with sensorimotor adaptation (Langan & Seidler, 2011; Schaffert et al., 2017) and motor sequence learning (Bo et al., 2009; Bo & Seidler, 2009; Chan et al., 2015) in studies that typically utilized the dominant hand. However, future work will include efforts to replicate the effects in the current study in other effectors.

We also acknowledge the high level of education in this sample, which may influence the nature or generalizability of our findings. And while the emergence of two principal components is interesting from a neuropsychological perspective by demonstrating that, at least in older adults, some visuospatial tests are similar to each other while others are very different and independent from one another, another limitation of this study is related to the clinical interpretability of the principal components themselves. For instance, tests of visual construction loaded onto the first (Block Design) and second (Figure Copy) principal components, suggesting that abstract-problem solving and visual memory/graphomotor function were the discerning differences between the principal components, respectively. Future work will disentangle if graphomotor function or visual memory (or both) explains why the Rey-Osterrieth Complex Figure test scores predict one-month motor learning outcomes.

While performance on many of these visuospatial tests involves executive function, it is important to note that two principal components emerged from our analysis, with PC2 primarily comprising the Rey-Osterrieth tests. It is likely that PC2 represents the psychological processes unique to the Rey-Osterrieth tests rather than a measure of executive cognition; otherwise, more visuospatial tests would have loaded on it. While we are unable to discern the latent constructs that PC1 and PC2 represent (and is outside the scope of the purpose of this study), it is clear that PC2 has a relationship with motor skill learning while PC1 does not. We acknowledge the principal component approach has limited clinical interpretability, yet it is necessary to control for statistical confounds and minimize Type-I error rate. For ease of clinical interpretation, however, a model comparison between individual mixed-effects models using each visuospatial test as a predictor of one-month motor learning outcomes is provided in the online-only Supplementary Material (eTable 2), which is consistent with the results of our main analyses.

It is also noted that the present study did not replicate previous correlations between the Benton Judgement of Line Orientation test and motor skill retention (Lingo VanGilder et al., 2018). A major distinction between these studies is the retention period over which the trained skill was retested (i.e., one week versus one month). A theoretical model of early-and-late stages of motor learning (Doyon & Benali, 2005) posits that skill improvement during early learning relies upon higher-order cognitive processes (e.g., visuospatial function), whereas improvement during later stages (e.g., automaticity has been achieved) shifts dependence from cognitive to specialized sensorimotor networks (such as cortico-striatal and -cerebellar circuits). Seminal work by Fleishman and Rich (1963) demonstrated this by showing that within a cohort of young adult males, the degree of motor skill improvement during early trials of a joystick-coordination task was correlated with visuospatial perception, whereas improvement during later stages of learning became less so (skill improvement during later trials were correlated with proprioceptive ability). Thus, the one-month retention period in the present study may indeed be a measure of late-stage motor learning in which the motor skill could be readily executed with negligible cognitive effort (Fitts & Posner, 1967). To further explore the reproducibility of previous findings, future work will evaluate if visuospatial perception and proprioception predict skill improvement during early and late stages of motor learning (i.e., one-week and one-month retention), respectively.

Due to its sensitivity and specificity to visuoconstructional deficits, the Rey-Osterrieth Complex Figure test has high clinical utility, such as diagnostic testing for i) numerous neurological pathologies like prodromal Alzheimer’s Disease (Han et al., 2015) and William’s syndrome (Hoeft et al., 2007; Mervis et al., 1999); ii) distinguishing dementias like Alzheimer’s and Parkinson’s Disease (Freeman et al., 2000; Possin et al., 2011); and iii) localizing nonverbal memory in patients with epilepsy (Frank & Landeira-Fernandez, 2008). Findings from the present study suggest that an older adult’s performance on the Rey-Osterrieth Complex Figure test is predictive of their motor learning capacity; future work will evaluate if this test may also be prognostic of an older adult’s capacity to benefit from motor rehabilitation, particularly when task-specific training is used as a therapeutic intervention. (In a sense, this is similar to the use of visuospatial and other cognitive tests to predict disease progression in more cognitively impaired adults, as summarized in Prado et al. (2019)). In other words, there is a clinical expectation that the learned motor skill will generalize to other motor tasks outside of therapy (i.e., skill transfer), thereby maximizing the benefit of training (Babulal, Foster, & Wolf, 2016). Thus, administration of the Rey-Osterrieth test prior to therapy may provide critical insight into an older adult’s ability to transfer motor skills learned during task-specific training. While administration of Delayed Recall is more time-intensive, results also indicate that these scores may uniquely estimate the durability of task-specific training (i.e., the amount of motor skill individuals learn and retain over a one-month period of no practice). To determine if the Rey-Osterrieth Complex Figure test can be used to predict motor rehabilitation response, future studies will evaluate if the observed behavioral relationship persists within a clinical population (e.g., stroke survivors) and the feasibility of implementing these tests to prognosticate an individual’s progression through motor rehabilitation.

### 4.3 Conclusions

Results of the present study indicated that among older adults, the degree of skill transferred one month after extensive upper extremity motor training may be explained, in part, by performance on the Rey-Osterrieth Complex Figure test. Results also suggest the neural pathways underlying visuospatial construction may be necessary for the generalizability of motor learning, rather than its durability in response to extensive training. Future studies will evaluate if this test can be administered prior to motor training to relate to risk of non-responsiveness in therapy and determine the structural neural correlates of this predictive relationship.

## Footnotes

1 That is, the major goal of this study was to see which of six clinically-available visuospatial measures is the strongest correlate of individual differences in the transfer of learned motor skills. No matter the outcome, the next logical conclusion is another experiment that is a better operationalization of the research question. Thus, a false negative potentially halts a fruitful area of work from moving forward, but a false positive would simply fail to replicate in the (necessary) subsequent study. As such, we reasoned that false negatives were more costly than false positives at this juncture.

