## Supplementary Data for "Evidence for associations between Rey-Osterrieth Complex Figure test and motor skill learning in older adults"

**Supplementary Material**

**Bivariate scatterplots**

The relationships between participant performance on each visuospatial test are shown in the bivariate scatterplot results below and indicates that specific visuospatial tests are linearly correlated, reflecting the principal component analysis results.

**eTable 1. Bivariate scatterplots of visuospatial tests.**

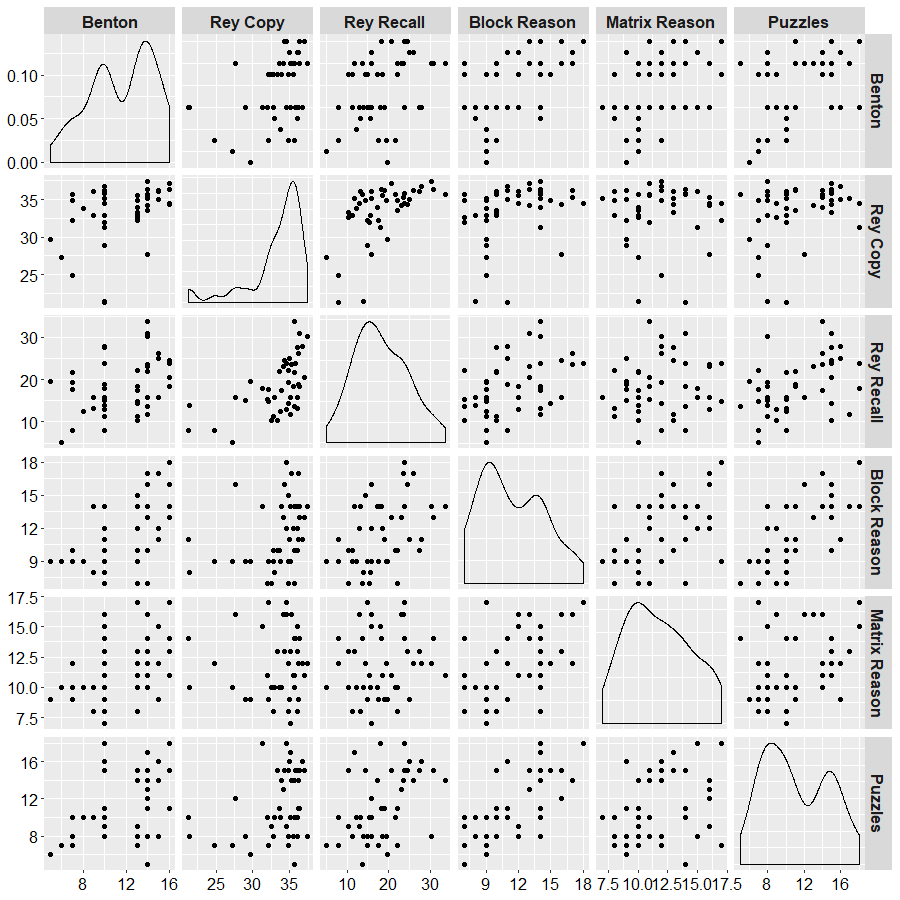

**Mixed-effect regression model comparison**

A separate mixed-effects model was used to evaluate each individual visuospatial test to avoid potential collinearity between predictor variables. The corrected Akaike’s Information Criterion (AICc) value of each regression model was used to evaluate each model’s goodness of fit, as it is an unbiased estimate of the long-run predictive deviance, where the model with the lowest AICc value explains the most variance and has the highest likelihood of replication in future samples. It is noted, however, that performing individual regression models could theoretically increase Type-I error. The AICc value of each regression model was used to determine each model’s fit (eTable 1). Consistent with results from the principal component analysis, the model including Rey-Osterrieth Complex Figure Copy had the lowest AICc value, followed closely by its Delayed Recall, suggesting that this test had the strongest relationship with one-month retention and transfer of motor training.

**eTable 2. Corrected Akaike’s Information Criterion value for each regression model.**

| **Visuospatial test** | **AICc** |
| --- | --- |
| Line Orientation | 609.89 |
| *Rey Figure Copy | 606.36 |
| Rey Delayed Recall | 607.27 |
| Visual Puzzles | 609.35 |
| Matrix Reasoning | 609.30 |
| Block Design | 609.98 |

AICc = corrected Akaike’s Information Criterion. *indicates model of best fit.
